# Evaluating the effect of denoising submillimeter auditory fMRI data with NORDIC

**DOI:** 10.1101/2024.01.24.577070

**Authors:** Lonike K. Faes, Agustin Lage-Castellanos, Giancarlo Valente, Zidan Yu, Martijn A. Cloos, Luca Vizioli, Steen Moeller, Essa Yacoub, Federico De Martino

## Abstract

Functional magnetic resonance imaging (fMRI) has emerged as an essential tool for exploring human brain function. Submillimeter fMRI, in particular, has emerged as a tool to study mesoscopic computations. The inherently low signal-to-noise ratio (SNR) at submillimeter resolutions warrants the use of denoising approaches tailored at reducing thermal noise – the dominant contributing noise component in high resolution fMRI. NORDIC PCA is one of such approaches, and has been benchmarked against other approaches in several applications. Here, we investigate the effects that two versions of NORDIC denoising have on auditory submillimeter data. As investigating auditory functional responses poses unique challenges, we anticipated that the benefit of this technique would be especially pronounced. Our results show that NORDIC denoising improves the detection sensitivity and the reliability of estimates in submillimeter auditory fMRI data. These effects can be explained by the reduction of the noise-induced signal variability. However, we also observed a reduction in the average response amplitude (percent signal), which may suggest that a small amount of signal was also removed. We conclude that, while evaluating the effects of the signal reduction induced by NORDIC may be necessary for each application, using NORDIC in high resolution auditory fMRI studies may be advantageous because of the large reduction in variability of the estimated responses.

## 1 Introduction

In recent years, the use of ultra-high field (UHF) magnetic resonance imaging has rapidly increased for a variety of applications. At UHF, the signal-to-noise ratio (SNR - Vaughan et al., 2001) and the blood-oxygenation-level-dependent (BOLD) contrast (Ogawa et al., 1992), the basis of functional MRI (fMRI), increase (Yacoub et al., 2001). This results in higher sensitivity to fMRI responses compared to more conventional field strengths (e.g. 3T and below). This allows for enormous benefits for the study of human brain function, in particular, the ability to acquire high spatial resolution (below 1 mm isotropic voxels) images. As such, at UHF it is possible to breach into the mesoscale and investigate fundamental computational structures and organizations of cortical functions, such as layers and columns (see e.g. De Martino et al., 2018; Dumoulin et al., 2018; Huber et al., 2015; Kok et al., 2016; Lawrence et al., 2019; Moerel et al., 2021; Olman et al., 2012; Uğurbil, 2018; Yacoub et al., 2008; Zimmermann et al., 2011).

The functional contrast to noise ratio (fCNR) in fMRI is dependent on the signal change compared to baseline and both physiological and thermal noise. Submillimeter fMRI at UHF trades the higher SNR for spatial resolution, often times leaving the resulting data in a thermal noise dominated regime (characterized as unstructured, zero-mean Gaussian distributed noise) emanating from electrical sources inherent to MRI hardware (Triantafyllou et al., 2005, 2011). This makes approaches oriented towards reducing thermal noise (i.e. improving the image SNR) of particular interest for neuroscience applications that require mesoscopic level imaging. Importantly, approaches for improving image SNR have to be evaluated against any practical considerations or tradeoffs, for example, their ability to preserve spatial information content at the finest scales (e.g. laminar and columnar cortical responses) (Polimeni et al., 2018) or whether any unwanted biases are introduced (Kay, 2022). Extensive averaging, one of the possible approaches for reducing thermal noise, could in principle help in highlighting small functional changes without altering the signal content. However, this approach – which assumes constant responses to the same stimuli over extended periods of time – is limited by practical implications such as the overall length of scanning sessions and the need for aligning data across multiple imaging sessions. While precision imaging approaches, that collect extensive data in only a few individuals, are becoming increasingly interesting in particular settings (see e.g. Allen et al., 2022; Michon et al., 2022; Poldrack et al., 2017) their application to mesoscopic imaging is far from standard and may not suffice when questions are oriented to generalizing effects at the population level. As an alternative to averaging, spatial smoothing could be used to increase image SNR. However, its application needs careful consideration as it comes with inevitable loss of spatial specificity (Turner & Geyer, 2014), which can be controlled if combined with anatomically informed constraints (e.g. laminar smoothing maintains specificity in the cortical depth direction while smoothing only tangentially) (Huber et al., 2021; Kiebel et al., 2000). Apart from averaging and (image) smoothing (with anatomical constraints), approaches for improving the detectability of effects (i.e. overcoming the limitations of low SNR regimes) have been considered at the analysis stage. Multivariate analyses, for example, have been argued to better leverage the information present in fine grained patterns and in part overcome the lower SNR of high resolution functional images, but may have some limitations in interpretability (Formisano & Kriegeskorte, 2012). In univariate analyses, the definition of noise regressors (through e.g. principal component analysis - Kay et al., 2013), has also been considered in order to improve the detectability of effects of interest, but relies on knowledge of the experimental design and assumptions such as the definition of noise pools (i.e. a collection of voxels whose time series is mostly representing noise sources). Denoising based on independent component analysis (ICA) has also been developed in fMRI and evaluated primarily in its ability to remove structured noise components (Pruim et al., 2015) and improving detectability of effects in lower resolution functional data that are mainly challenged by physiological noise (Griffanti et al., 2014; Salimi-Khorshidi et al., 2014). For completeness it is important to note that approaches to remove structured (physiological) noise in fMRI (and thus not tailored to the reduction of thermal noise) include, apart from ICA, the use of multiple echoes to estimate sources of variance (Gonzalez-Castillo et al., 2016; Steel et al., 2022), or measuring physiological data to subsequently remove the noise sources from the data (e.g. RETROICOR - Glover et al., 2000; or RETROKCOR - Hu et al., 1995).

A denoising technique tailored to the removal of thermal noise that has recently been introduced is NOise Reduction with DIstribution Corrected Principal Component Analysis (NORDIC PCA - Moeller et al., 2021; Vizioli et al., 2021). NORDIC is a pre-processing approach based on PCA that selectively removes components that are indistinguishable from zero-mean normally distributed noise (see e.g. https://layerfmri.com/2023/07/10/nordic/#more-3956 for an informal description of the approach). Compared to other PCA denoising techniques (see Veraart et al., 2016), the main difference rests in the approach used to estimate the number of (principal) components that are removed (i.e. the threshold on the eigenvalue spectrum that distinguishes noise components from signal components). NORDIC has been initially extensively evaluated on visual cortical responses elicited by blocked (temporally prolonged ∼ 12 seconds) stimulation and has been shown to increase detection sensitivity without affecting the overall signal change and spatial precision of the responses (i.e. without introducing spatial blurring - Vizioli et al., 2021). NORDIC has also been evaluated and compared to other PCA based denoising approaches (dwidenoise - Cordero-Grande et al., 2019; Manzano-Patron et al., 2023; Veraart et al., 2016). Compared to dwidenoise and more conventional smoothing approaches, and in experimental designs ranging from blocked to event related visual stimulation, NORDIC has been shown to better preserve local and global spatial smoothness of the functional data as well as the temporal characteristics of the responses (i.e. temporal smoothing) and it has been shown to not introduce unwanted effects (Dowdle et al., 2023; but see Fernandes et al., 2023 for an evaluation in rodent data). NORDIC has been rapidly picked up by the community and its usability is now being examined across different areas (including visual and motor regions), field strengths (3T and 7T), and acquisition techniques (see e.g. Dowdle et al., 2022; Knudsen et al., 2023; Raimondo et al., 2023). These recent studies consistently show that NORDIC improves detectability of the effects. However, while NORDIC has been shown to improve (statistical) signal detection, generalizing these results to other cortical regions and to designs that are particularly SNR limited (e.g. auditory cortical responses elicited by slow event-related designs) still requires careful evaluation of its benefits as opposed to any potential unwanted bias.

Here we focus on the application of NORDIC to submillimeter functional MRI data collected to investigate auditory cortical responses elicited by a slow event-related design. The auditory cortex is located next to large air cavities, with parts of it like primary cortical regions lying further away from the receive coils compared to other sensory regions (e.g. visual and somatosensory regions). These and other factors (e.g. the need for large field of views to image bilateral auditory cortical areas) make imaging auditory cortical regions sensitive to geometric distortions and signal dropouts due to large B_0_ inhomogeneities (Moerel et al., 2021) and not only for BOLD type acquisitions (Faes et al., 2023). Furthermore, the percent signal change elicited in auditory regions is lower than in visual cortex (De Martino et al., 2015). However, despite these challenges, there have been several high resolution auditory studies that look at cortical depth dependent responses (see e.g. Ahveninen et al., 2016; De Martino et al., 2015; Gau et al., 2020; Moerel et al., 2018). As such, given that auditory submillimeter studies are especially restricted by low SNR, they would greatly benefit from thermal noise reduction. However, the efficacy of PCA-based denoising methods also depends on the relative contribution of signal and noise. Therefore, submillimeter auditory fMRI may present a challenge for NORDIC. Collectively, these considerations warrant the need to explore the effect of NORDIC denoising on submillimeter fMRI data collected in the auditory cortex. We center our evaluation on the improvements in tSNR by considering changes to both the mean percent signal change and its variability.

## 2 Methods

### 2.1 NORDIC

NORDIC is a denoising approach that operates on either complex-valued or magnitude-only fMRI time series. As the use of parallel imaging results in a spatially varying amplification of the thermal noise according to the g-factor (Pruessmann et al., 1999), if necessary, the NORDIC algorithm first normalizes the functional data by the g-factor, resulting in the thermal noise being uniformly distributed across space (to fulfill the assumption of PCA denoising that noise is identically distributed across voxels). NORDIC uses a locally low rank approach to perform a patch-wise PCA across space and time. An estimate of the noise level is obtained from an appended acquisition without a radiofrequency excitation (e.g. a noise scan) or an estimate of the g-factor noise (Ma et al., 2020). In each patch, the noise threshold defines the principal components that are removed from the eigenspectrum as they are considered to be indistinguishable from zero-mean Gaussian distributed noise. The noise threshold is chosen with Monte-Carlo simulations for a Casorati matrix with zero-mean normally distributed sampling and, depending on the settings, considering an ideal or realistic noise distribution. After the removal of noisy principal components, the patches are recombined and the g-factor is re-applied to reconstruct the fMRI images. Assuming signal redundancy within the patch (i.e. enough voxels carrying the same information), NORDIC aims at removing thermal noise from the time series while preserving the fine-grained temporal and spatial structure of the signal that is assumed to be carried by the preserved principal components. For more details on NORDIC we refer to the original publications (Moeller et al., 2021; Vizioli et al., 2021).

Currently, there are two implementations of NORDIC available (https://github.com/SteenMoeller/NORDIC_Raw). We focus on the use of (NIFTI_NORDIC - version 04-22-2021) which takes nifti formatted data of both magnitude and phase images as input.

### 2.2 MR imaging acquisition

Data was collected with a 7T Siemens Magnetom System with a single channel transmit and 32-channel receive NOVA head coil (Siemens Medical Systems, Erlangen). Whole-brain anatomical T1-weighted images were collected using a Magnetisation Prepared 2 Rapid Acquisition Gradient Echo (MP2RAGE) sequence at a resolution of 0.75 mm isotropic (192 slices, TR = 4300 ms, TE = 2.27 ms) (Marques et al., 2010).

Functional data were acquired with 2D gradient-echo (GE) echo planar imaging (EPI) along with simultaneous multi-slice (SMS)/(MB) multiband (Moeller et al., 2010; Setsompop et al., 2012) (0.8 mm isotropic, 42 slices, TR = 1600 ms, TE = 26.4 ms, MB factor 2, iPAT factor 3, 6/8 Partial Fourier, bandwidth 1190 Hz, field of view: 170 x 170 mm, matrix size: 212 x 212, phase encoding = anterior to posterior; coil combination = SENSE1).

### 2.3 Participants

Ten healthy participants took part in this fMRI study (aged between 23 and 69 years old, 5 females). Participants had no history of neurological disease or hearing disorders. Eight participants were scanned at the Center for Magnetic Resonance Research in Minneapolis (CMRR) and two were scanned at New York University (NYU) using the identical imaging protocol except for slight differences in TR (TR_CMRR_ = 1600 ms, TR_NYU_ = 1650 ms). The local IRB at the individual institutions approved the experiment. All participants signed informed consent forms before commencing the study.

### 2.4 Experimental design

Participants passively listened to tone sequences. Conditions were designed to investigate predictive processing in the auditory cortex (based on sequences used in Berlot et al., 2018), but we will disregard the neuroscientific purpose of the experimental paradigm and focus on the effect of denoising instead.

Six conditions were presented. All conditions consisted of sequences of four tones. The four tones were presented for 100 ms each with a 400 ms gap between tones (total tone sequence length was 1.6 seconds). The conditions were designed such that the first 3 tones were ‘contextual’ tones ordered in either a descending, ascending or scrambled fashion. The frequencies used for these contextual tones were always the same three (493.9, 659.3 and 987.8 Hz), albeit presented in a different order. The fourth tone was selected such that three conditions ended in a high frequency (1318.5 Hz) and three conditions ended in a low frequency (329.6 Hz). This resulted in two predictable sequences (PredH and PredL), two mispredicted sequences (MispredH and MispredL), and two unpredictable sequences (UnpredH and UnpredL) as displayed in Figure 1. The auditory stimuli were presented concomitantly with the scanner noise (i.e. no silent gap for sound presentation was used).

**Figure 1.**
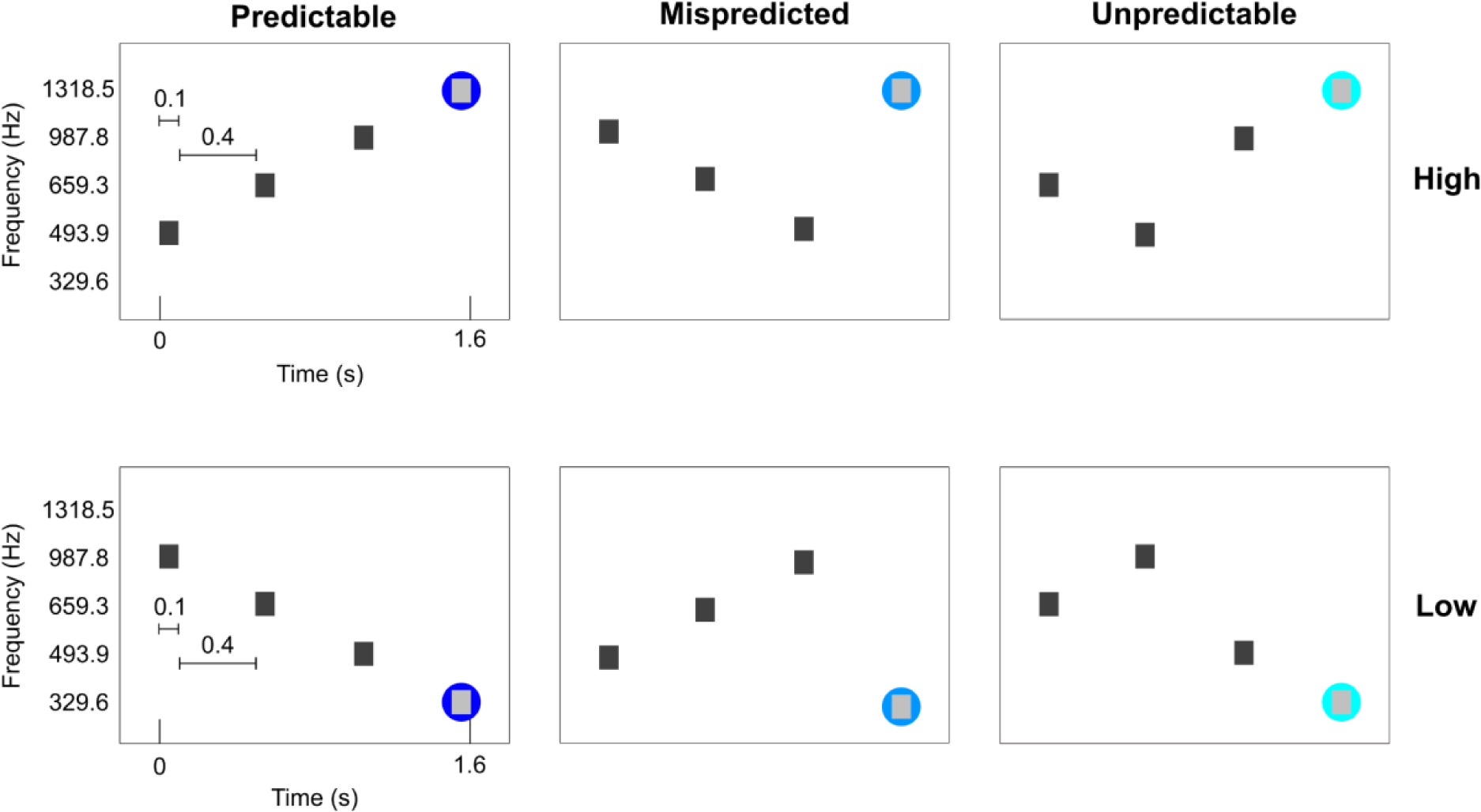
Experimental conditions. The first three tones are contextual eliciting a strong or weak prediction. The three contextual tones are presented at the same frequencies, albeit in different orders. The fourth tone can either be a high or low target frequency. The fourth tone can either consecutively follow the ascending or descending order (PredH and PredL), or the contextual tones could be deviant (MispredH and MispredL) or the contextual tones could be scrambled (UnpredH and UnpredL). The two predictable sequences were presented ten times per run, whereas the other four conditions were presented four times per run.

Per run, each of the predictable sequences were presented 10 times, the mispredicted sequences and the unpredictable sequences were each presented 4 times in a randomized order (for a total of 36 trials in one run). Tone sequences were presented in a slow event-related design with an average inter-trial interval of 6 TR’s (ranging between 5 and 7 TR’s). For each participant, we collected 6 to 8 runs that lasted approximately 6 minutes each (including noise scans at the end of each run). Magnitude and phase Dicom images were exported from the scanner.

### 2.5. Data preprocessing

#### 2.5.1 NORDIC preprocessing

Dicom files were converted to NIfTI format (separately for magnitude and phase images - MRIcron, version 1.0.2). The magnitude and phase NIfTI files were used as the input to NIFTI_NORDIC. We used two different settings for NORDIC denoising: 1) using default settings for fMRI data (PCA kernel size of 11:1, temporal phase = 1, phase filter width = 10, the noise scan is used for empirical noise estimation) and 2) the same as the default except for the use of the noise scan in the estimation of the noise threshold. In the NIFTI_NORDIC implementation, not using the noise scan results in using a noise threshold based on the g-factor estimation (in the implementation, a threshold of 1/sqrt(2)), which is generally more conservative (i.e. resulting in the removal of less principal components) than the estimated threshold when using the noise scan, reflecting the empirical observation that the approach for g-factor estimation underestimates the value by up to 10%. This resulted in three datasets (per run), the first, which will be referred to as the *‘Original’*, represents the fMRI data without NORDIC denoising. The second, which we will refer to as *‘NORDIC default’* (NORdef), represents the fMRI time series resulting from the processing with default NORDIC settings. The third, we will refer to as ‘*NORDIC No Noise*’ (NORnn), represents the fMRI time series resulting from the use of NORDIC without separate noise scans for the estimation of the noise threshold.

#### 2.5.2 Pre-processing

The anatomical and functional data were analyzed using a BrainVoyager software package (BV - version 21.4, Brain Innovation, Maastricht, The Netherlands) and custom Matlab scripts (The MATHWORKS Inc., Natick, MA, USA). After the initial NORDIC denoising step, functional processing was performed identically across datasets. The noise acquisitions were removed from each time series. Pre-processing of the functional data included slice scan time correction using sinc interpolation and motion correction along three dimensions using intrasession alignment to the run closest in time to the collection of opposite phase encoding images (run 1 in most participants, run 4 in two participants). In addition, temporal filtering was applied to remove low frequencies (high-pass filtering with 7 cycles per run) and high frequencies (temporal gaussian smoothing with a full width half maximum kernel of 2 data points). Reversed phase polarity acquisitions were used to correct for geometric distortions using Topup (FSL version 6.0.4). In one participant we experienced issues collecting opposed phase polarity images and therefore no distortion correction was performed in this participant.

The anatomical data were upsampled to 0.4 mm isotropic, corrected for inhomogeneities and transformed to ACPC space. A segmentation was created using the deep neural network in BV to determine the initial white matter (WM) and gray matter (GM) boundary and GM/cerebral spinal fluid (CSF) border. The segmentation of the temporal lobe was manually corrected in ITK snap (Yushkevich et al., 2006). With this corrected segmentation, we created mid-GM surface meshes in BV. Additionally, we estimate the cortical thickness of the high-resolution segmentation.

#### 2.5.3 ROI definition

Five bilateral regions of interest (ROIs) were drawn on the individual mid-GM meshes based on macro-anatomical landmarks (as described in Kim et al., 2000), covering the temporal lobe including Heschl’s Gyrus (HG), Planum Polare (PP), Planum Temporale (PT), anterior superior temporal gyrus (aSTG) and posterior superior temporal gyrus (pSTG). These ROIs were projected back onto the anatomy in volume space (extending 3 mm inwards and outwards from the mid-GM surface). These masks were first intersected with the GM definition and then dilated (six steps) in order to obtain the final masks that include GM as well as the WM and the CSF surrounding it. The union of all the masks (temporal lobe mask) was used to run the statistical analysis (General Linear Model, see below), while results were inspected separately per ROI in some analyses.

### 2.6. Analyses

#### 2.6.1. General Linear Model

All statistical analyses were performed with custom Matlab scripts. Time series were first normalized to percent signal change (PSC). For our first-level analysis, we fitted a general linear model (GLM) with single trials per condition as predictors (36 trials and one constant per run). Predictors were convolved with a standard two-gamma hemodynamic response function (HRF) that peaked at 5 seconds after the onset of the stimuli. In order to evaluate the effects that NORDIC has on the reliability of the responses, we obtained response estimates (beta weights) and computed statistical activation maps by considering the variability across single trials (i.e. beta time series) for all predictors combined (sounds versus no sounds) and for each condition separately. In other words, we here estimate the variance of the response considering the variability across trials and not the variance of the residuals of the GLM fit. This helped us in evaluating measures of reliability of the response estimates.

After the GLM, in each individual’s anatomical ROI (considering all voxels in the ROI) we evaluated: 1) the change in beta (PSC) per condition before and after NORDIC processing; 2) the change in single trial t-statistics (mean divided by variance across trials); 3) the spatial replicability of the mean betas (PSC); and 4) the spatial replicability of the t-statistics. For all individual subject data, all analyses were performed by randomly sampling half of the runs (i.e. repeated split half with 50 repetitions - Valente et al., 2021). The spatial replicability of the betas and t-statistics was computed by correlating the variable of interest (PSC or t-statistics) across the two random splits of the data. Finally, across all ROIs we investigated changes in beta values (before and after NORDIC) in relation to the tSNR. Note that we compute tSNR (defined as the mean divided by the standard deviation of the time series) on the original data after pre-processing (tSNR_pr_). This choice inflates the tSNR we report compared to the more conventional choice to calculate tSNR on the un-preprocessed data (in analyses not shown we confirmed that the results we report here are not dependent on the choice or pre-processing applied to the time series).

At the group level, interactions were tested with repeated measures ANOVA (where processing strategy is the repeated measure). Main effects were tested for significance using permutation testing by permuting, for each test, individual subject data across processing strategies (all possible permutations [2^10^]) and corrected for multiple comparisons using Bonferroni.

#### 2.6.2. Correlation and cross-validation analyses

To evaluate the spatial similarity of beta estimates across processing strategies we computed the correlation of the estimated beta maps. In particular, we considered: 1) the correlation of each NORDIC processed run (NORdef and NORnn) to the corresponding original run (separately for each of the six conditions); 2) the run-to-run correlation within each processing strategy (i.e. within Original, NORdef and NORnn data) and 3) using leave-one-run-out, the correlation of each run (i.e. run 3) to the average of all other runs (all runs except run 3). Importantly for this last analysis the reference model (i.e. the averaged map coming from all runs except one) was always kept to be the one extracted from the original time series.

At the group level, interactions (e.g. processing strategy and condition in the first analysis) were tested with repeated measures ANOVA (where processing strategy is the repeated measure). To do this, data where Fisher z-transformed prior to the ANOVA. Main effects were tested for significance using permutation testing by permuting, for each test, individual subject data across processing strategies (all possible permutations [2^10^]) and corrected for multiple comparisons using Bonferroni.

#### 2.6.3 Tonotopic maps

From the two predictable conditions we create tonotopic maps (as best frequency maps, see Formisano et al., 2003; Heynckes et al., 2023 for an exampe where the procedure is applied with only two frequencies, as is the case here). Tonotopic maps were computed in volume space and interpolated to the mid cortical surface.

#### 2.6.4 Variance Partitioning

We reasoned that the total variance from the original (magnitude) time series (per voxel) could be partitioned as follows:

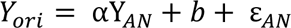

Where *Y_ori_* and *Y_AN_* are the original time series and the time series after NORDIC preprocessing respectively (*Y_AN_* can then come from either NORdef or NORnn). The values for the scaling factor and intercept were estimated with ordinary least squares (OLS), thus obtaining an estimate of the scaling and intercept (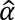 and 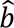). The estimated scaling and intercept are then used to compute (per voxel) estimated residuals 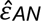. These residuals can be interpreted as the portion of the original time series that is orthogonal to the NORDIC time series. We refer to this as the residuals of the original time series after NORDIC (residuals after NORDIC in short). This decomposition guarantees that the total sum of squares of the original data (representing the variability in the data with respect to their mean) can be expressed as the sum of squares of the data after NORDIC (weighted by 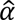) and the sum of squares of portion of the original data that is orthogonal to the data processed with NORDIC (i.e. the residuals after NORDIC 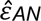). That is:

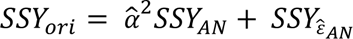

To quantify the variance associated with the experimental design in the original data, as well as the data after NORDIC processing and the residuals after NORDIC (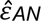), we regressed *Y_ori_*, *Y_AN_*, and 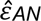 against our design matrix (X). This second regression allowed us to partition the variance that, in each of the three signals of interest (Y*_ori_*, *Y_AN_*, and 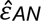), is related to the design 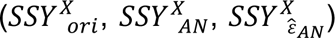, along with an error term for each.

We present the results by calculating the ratio of the sum of squares. First, within each processing strategy (Original, NORdef and NORnn), we compared the variance explained by the design to the total variance of each respective time series. Second, for the NORDIC processed data (NORdef and NORnn) we compared the variance associated with the design, in their respective residuals after NORDIC 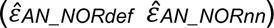, to the total sum of squares of the original time series. This last analysis allowed us to reveal the portion of the variance associated with the design that is not present in the NORDIC processed data and is thus removed by NORDIC.

#### 2.6.5 Laminar analysis

We explored the effect of NORDIC denoising on the cortical depth dependent estimates. Beta maps were computed across 11 cortical depths and sampled on the mid-GM surface in BV. These maps were subsequently intersected with a mask of HG. The single trial betas were averaged across vertices and subsequently across trials. The variability was computed across trials.

## 3 Results

### 3.1 Activation and spatial patterns

We assessed the effect of NORDIC on detection sensitivity by evaluating the overall activation (sounds > no sounds) elicited by single trials in our experimental design. Statistical maps were computed by considering the mean and variability (t statistic) across (single) trials (not the GLM residuals) and corrected for multiple comparisons using false discovery rate of qFDR<0.01. This is a more stringent threshold than the customary qFDR<0.05 because it allows better appreciation of the differences between processing strategies in each individual. Figure 2 presents the results in one exemplary volunteer (all other volunteers showed similar results - data not shown) on a representative transversal anatomical slice, highlighting the statistical advantage in detection sensitivity conferred by denoising. At the same statistical threshold both NORdef and NORnn resulted in more activation. In this volunteer, for example, 34% of voxels in our temporal lobe mask were significantly active at the qFDR threshold, whereas NORdef and NORnn resulted in 51% and 44% of voxels active, respectively. NORDIC denoising results in overall higher t-statistics, the 90th percentile across voxels for each of three datasets was 10.38, 13.42, and 12.38, respectively. These results are in line with previous applications of NORDIC (Dowdle et al., 2022, 2023; Knudsen et al., 2023; Raimondo et al., 2023; Vizioli et al., 2021) and similar to these previous reports, activation maps do not appear spatially distorted (i.e. blurred) when comparing NORDIC processed data to the original.

**Figure 2.**
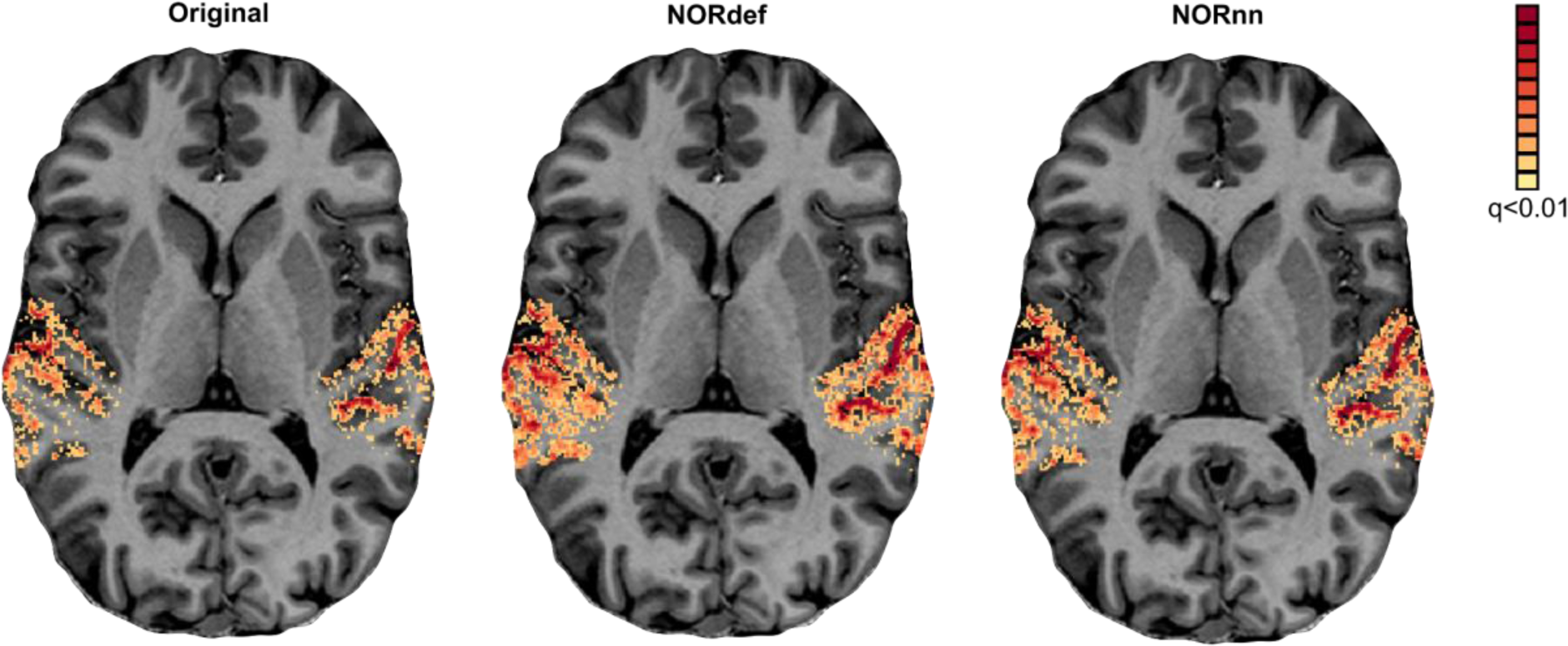
Single subject overall response to sounds (qFDR<0.01). From left to right we show the t-maps resulting from a GLM with single trials as predictors of the Original, NORdef and NORnn data on a transversal slice.

To evaluate some of these effects further, we analyzed the spatial patterns of activation (separately per condition). Figure 3A shows the similarity (correlation) of beta maps averaged across trials of NORdef and NORnn to the beta maps obtained from the original data (median and interquartile range across runs). NORnn resulted in a higher correlation to the original data compared to NORdef but the correlation values were similar across conditions. That is, the similarity was not influenced by the different amount of repetitions of specific conditions (e.g. the mispredicted and unpredictable conditions). In what follows, we present results of the predictable condition(s) only. Figure 3B shows the run-to-run reproducibility of the spatial patterns of activation within each processing strategy for PredH. NORdef and NORnn resulted in more reproducible spatial patterns compared to the original dataset. These first two analyses show that NORdef and NORnn effectively reduce thermal noise and improve reliability of the estimates (Figure 3B), while NORnn preserves more similarity to the original data. In the absence of a ground truth, we reasoned that the spatial pattern elicited by averaging multiple runs of the original data would be a reasonable choice to compare the results of single runs in their ability to approximate results obtained with higher SNR. To this end, we computed the average of the spatial pattern of activation elicited by PredH in the original data in all but one run. This reference pattern was correlated to the left out run in the original data and to the same run after NORDIC processing. We repeated this analysis each time leaving a different run out. The results (Figure 3C) show that after NORDIC, activation patterns in individual runs are more similar to the reference.

**Figure 3.**
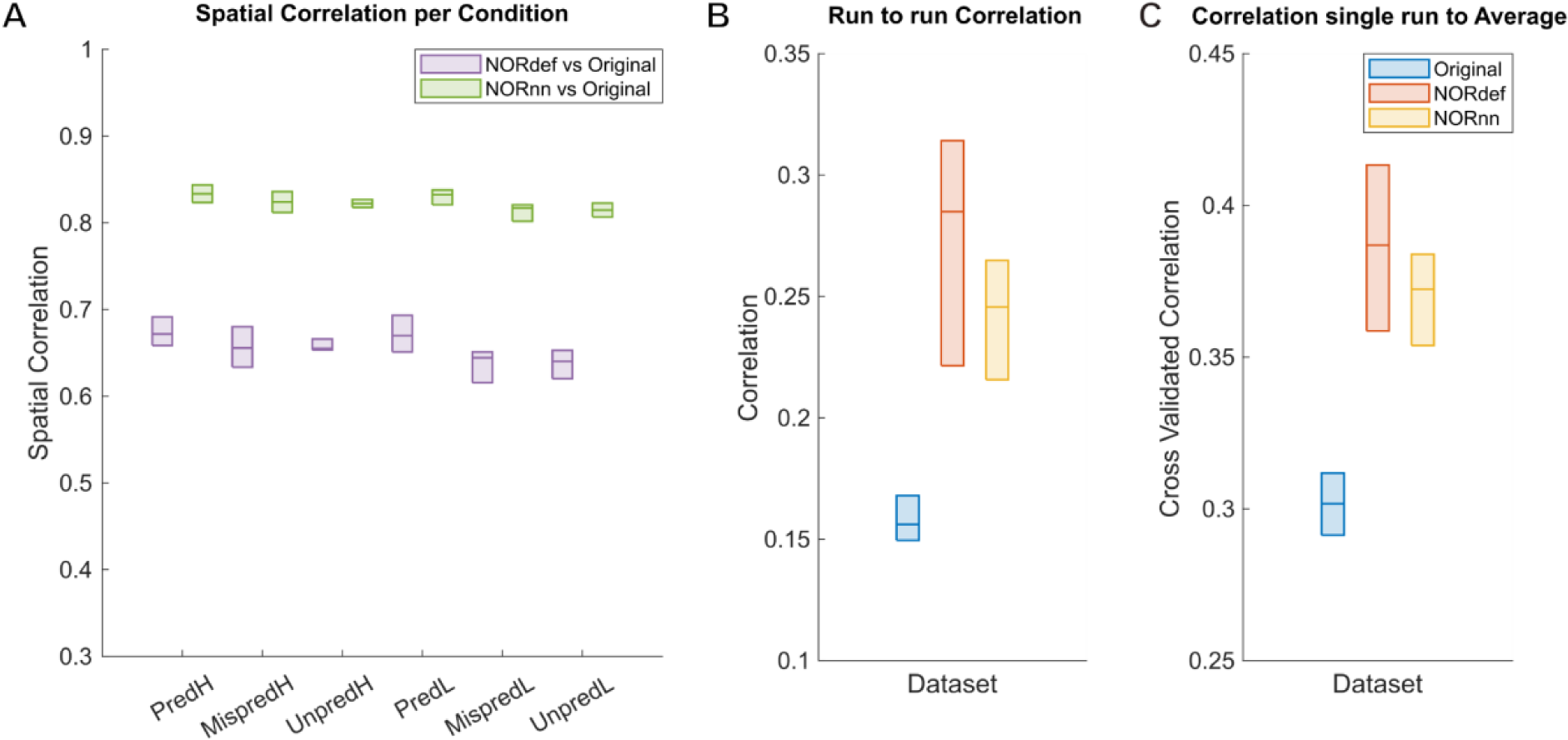
Single participant correlation analyses. Box charts display the median and interquartile ranges. A) Spatial correlations of beta maps for each condition. There is no difference in correlations between conditions. The correlation values between NORnn and the Original dataset are higher, indicating that noise removal in NORnn is more conservative than in the NORdef dataset. B) Run-to-run pairwise correlations computed per dataset for the PredH condition. Beta estimates across runs become more similar in both denoised datasets, albeit the estimates are more stable in the denoised data. C) Cross-validated correlation of one run to the average of n-1 runs of the Original data for the PredH condition. Both denoised datasets are more similar to the average of the Original dataset.

Figure 3 reports the result in a representative volunteer, while the group results (median and interquartile range across all our volunteers) is presented in Figure 4 (considering the variability across the mean estimates of every subject). The group results support the trend seen in the single-subject analysis, except in two individuals that showed very little improvement in either run-to-run variability or correlation to the reference pattern obtained in the original data (light gray dots in Figure 4B and C). Correlation coefficients were compared with a two-way repeated measures ANOVA (with condition and processing strategy as factors). There was no interaction between condition and processing strategy. Permutation testing showed a main effect of method, (p<0.001) indicating that at the group level NORnn results in a larger similarity of the spatial patterns to the original data. This is in line NORnn being more conservative, that is, resembling more the original data (due to a lower noise threshold and the removal of less noise components). The stability of run-to-run estimates (Figure 4B) was significantly higher for the NORnn compared to the Original data (p=0.041), whereas there was no evidence of a difference between NORdef and the Original data (p=0.064). At the group level, the correlation of a single run to the average of our reference was not significant in either NORdef compared to the Original data (p=0.258) or NORnn compared to the Original data (p=0.053).

**Figure 4.**
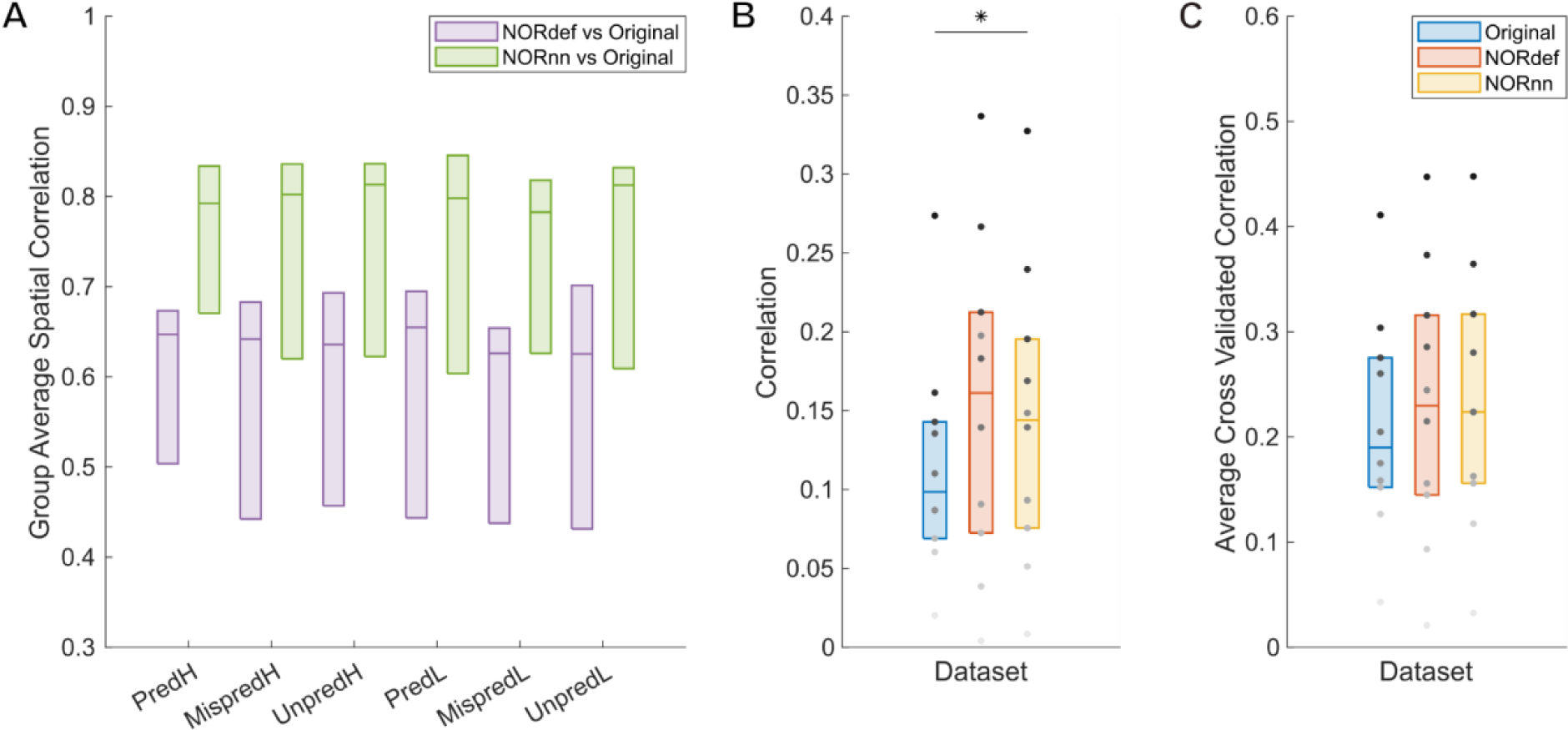
Group Figure of the same analysis as Figure 3. A) Across conditions, correlations between NORDIC denoised datasets and the Original data are indistinguishable indicating that number of repetitions do not affect the effect of NORDIC denoising. B) In general, stability of beta estimates increases with the use of NORDIC denoising. Gray dots indicate different participants. C) Average cross validated correlation values of single runs to the average of the Original data, for both predictable conditions. * indicates p<0.05, ** indicates p<0.01.

Our design also allows the derivation of tonotopic maps, albeit from only two frequencies, by computing best frequency maps on the predictable high and predictable low conditions. Figure 5 shows, for a single left hemisphere, tonotopic maps projected on the mid-GM surface intersected with their respective t-map. As expected (Moerel et al., 2014), a low frequency preferring region is visible along Heshl’s gyrus (HG) surrounded by two high frequency preferring areas. This gradient is visible in the Original data and becomes more discernible in the NORnn and NORdef tonotopic maps, respectively. The fact that some regions are more clearly preferring one of the two frequencies (i.e. blue regions anterior to HG) highlights the fact that after NORDIC the frequency preference is more spatially homogeneous and less corrupted by noise.

**Figure 5.**
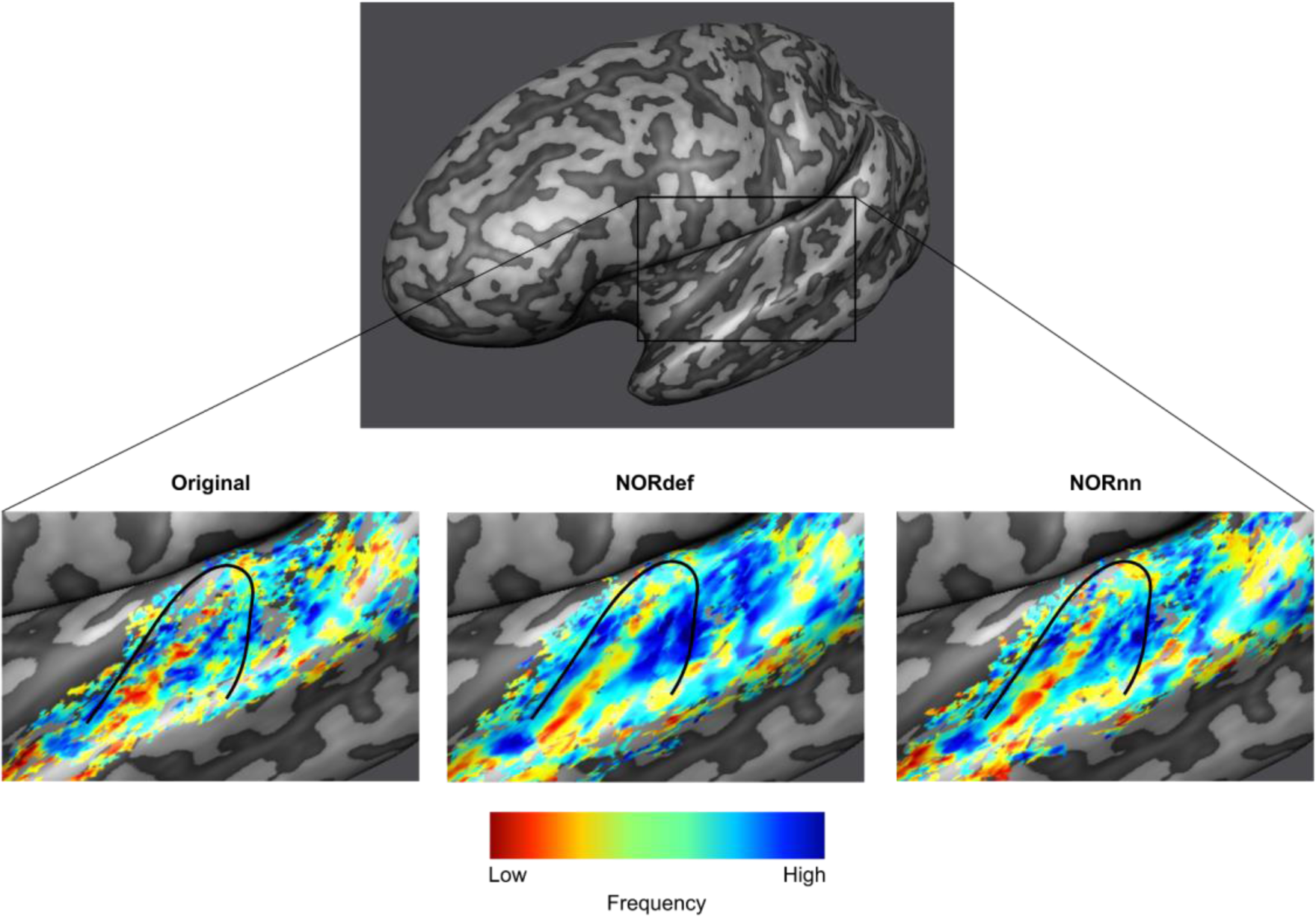
Tonotopic maps. Frequency preference maps are computed for each dataset and for one example participant we display these maps on an inflated mid-GM surface. Denoising does not seem to alter the frequency preference as the high-low high gradient is visible in all three datasets. The maps computed from the denoised datasets are less noisy.

The previous applications of NORDIC (Dowdle et al., 2023; Vizioli et al., 2021) have shown that detection sensitivity with NORDIC comes due to a reduction in variance without any change to the percent signal response. While this effect would explain our results at the level of the whole temporal lobe (reported in Figures 3 and 4), we investigated changes in percent signal as well as its variability across trials also in separate anatomically defined ROIs. In the temporal lobe, across all ROIs, NORDIC denoising resulted in reduced percent signal change (Figure 6A). This reduction was more pronounced in the NORdef compared to NORnn. Changes in PSC though come with a larger change in variability of the response when using NORDIC. This is clear when considering t-values within each of the ROIs (Figure 6B). The increase in t-values is most apparent in the NORdef time series. These changes induced by NORDIC processing are visible in ROIs that are activated by our design (i.e. the pattern is less visible in the aSTG that has little activation in our experiment). The change in betas induced by NORDIC is most evident in voxels whose overall signal level is low (see Figure 7). The bias introduced by NORDIC in the single ROIs does not come with detrimental effects to the reliability of the estimates in each ROI compared to the analysis at the level of the whole temporal lobe. When analyzing the reliability of spatial patterns in the individual ROIs (Figure 6C and D for PSC and t-value, respectively) the results are in line with the previously reported pattern at the level of the whole temporal cortex (Figures 3 and 4), that is NORDIC processing is associated with a general improvement in reliability.

**Figure 6.**
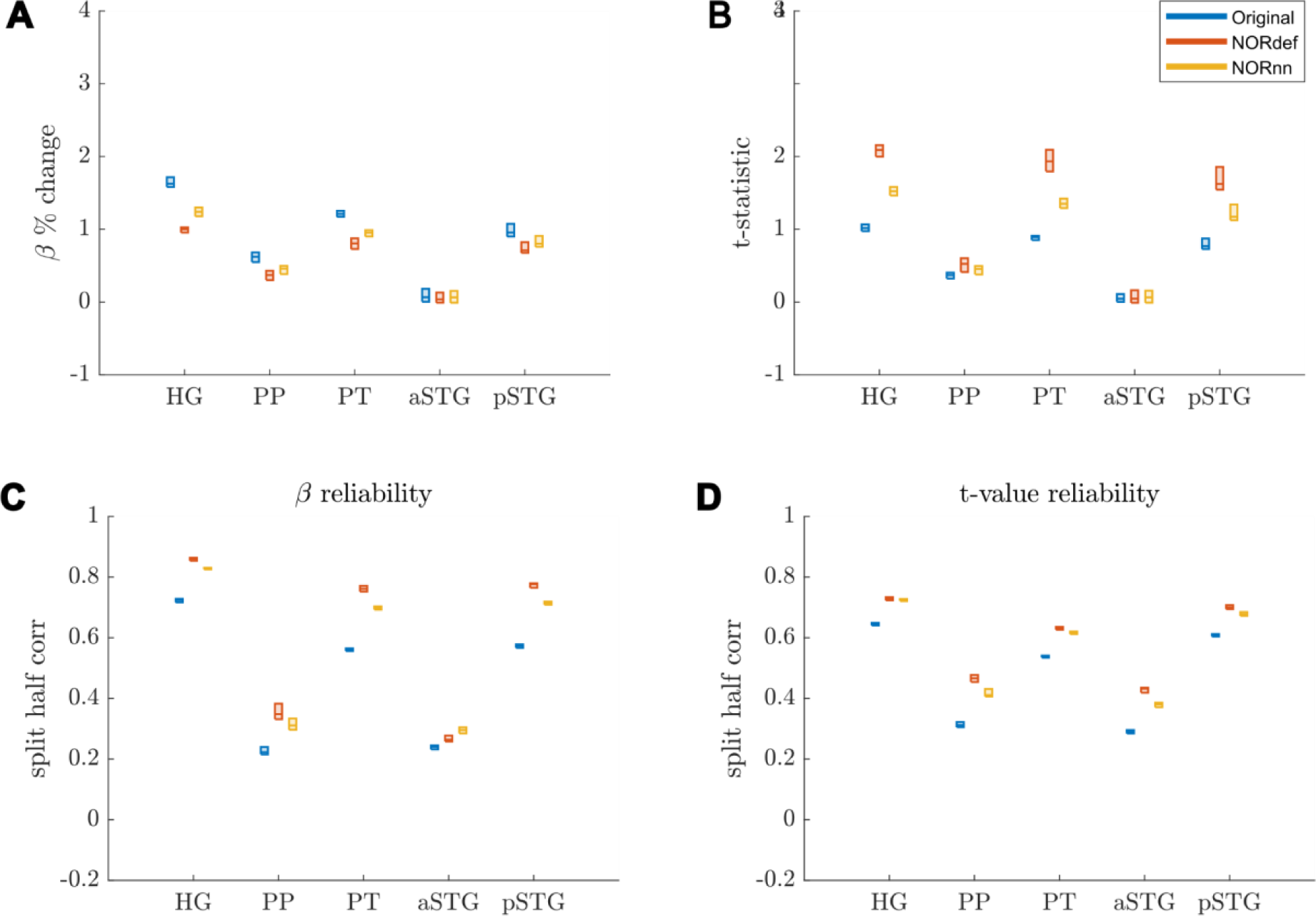
Responses in gray matter confined to regions of interest. A) Beta values calculated in percent signal change. In each ROI where there is signal present in the Original dataset, we observed a reduction in beta values after denoising. This reduction was lower in NORnn. B) T-statistics are increased after denoising, which was most pronounced in the NORdef dataset. C) Split half correlations were calculated to estimate the stability of beta responses. This revealed that beta values are more stably estimated after denoising. D) T-values are more reliably estimated in NORDIC.

**Figure 7.**
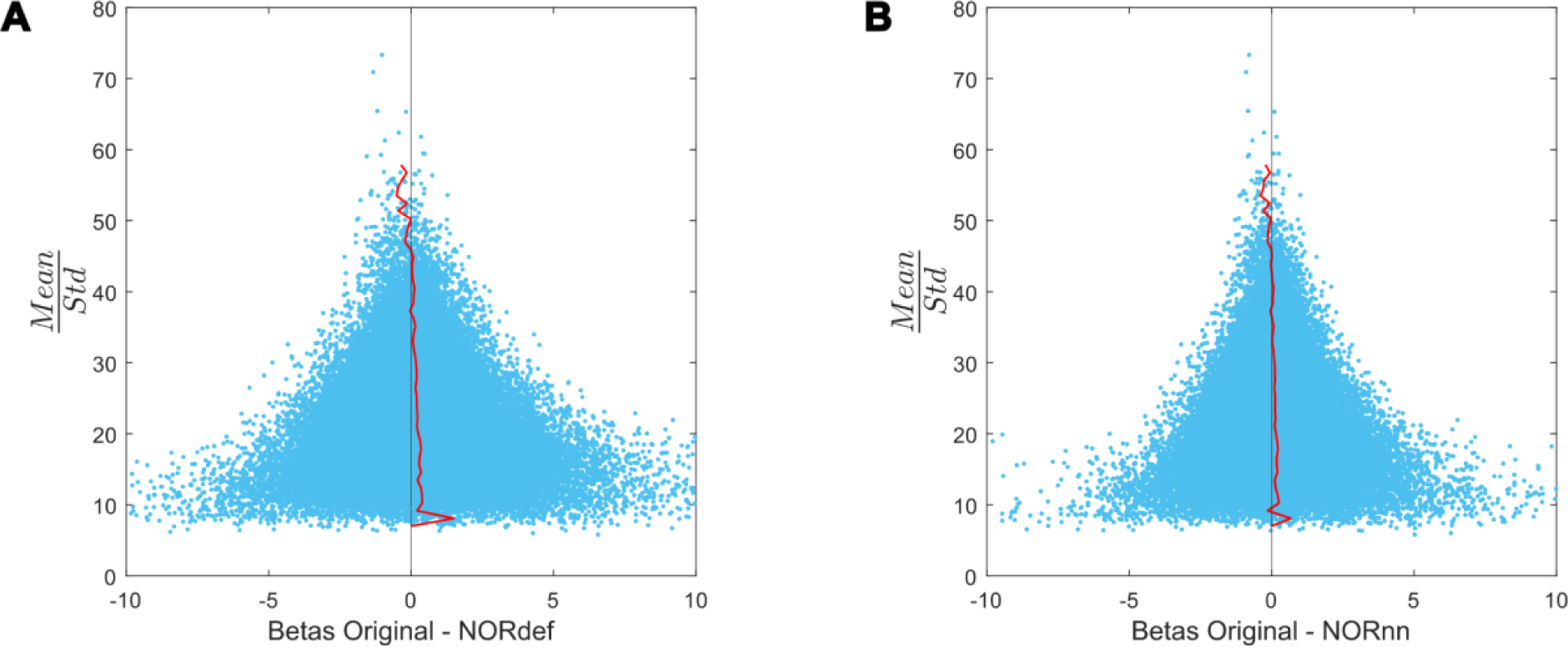
Beta difference in relation to tSNR for one representative subject. A) Betas before and after NORDIC are displayed as a function of mean/standard deviation (tSNRpr). For low tSNRpr values, the betas change in both directions. However, at high tSNRpr, the betas remain relatively similar after NORDIC. The red line indicates the mean beta difference per bin. The black line indicates a beta difference of zero. B) Same as A but for the beta difference between Original minus NORnn betas.

At the group level (Figure 8), a similar result becomes apparent. These results indicate that, in our data, there is evidence for a bias-variance tradeoff associated with the application of NORDIC. Repeated measures ANOVAs showed a significant interaction between processing strategy and ROI for each of the subfigures of Figure 8 (all p-values were smaller than 0.001). Per ROI we subsequently tested all three comparisons using permutation testing and corrected for multiple comparisons. The resulting p-values can be found in Table 1 in the Supplementary Materials.

**Figure 8.**
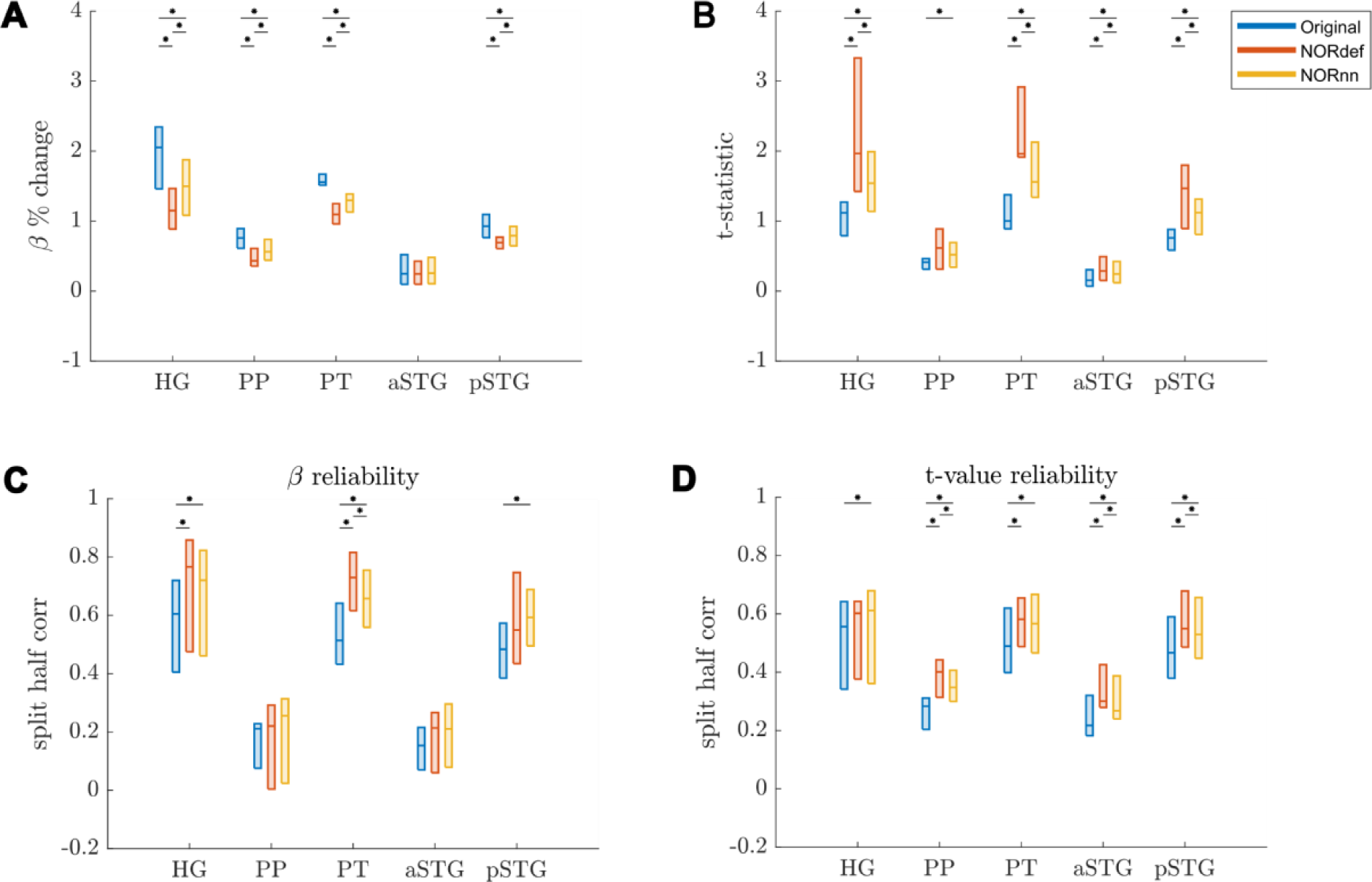
Group Figure of beta- and t-value estimates. A) Average reduction of beta values across participants. B) At the group level, the increase in t-values remains. C) On average, denoising results in a better estimate of beta values calculated with split half correlations in ROIs where there is more signal in the data. D) t-value reliability is generally higher after NORDIC than in the Original data. * indicates p<0.05.

### 3.2 Variance Explained

To investigate the nature of the bias introduced by NORDIC further, we quantified the variance explained by the design both in the time series as well as in the portions of the original time series that are not present in either the NORdef or NORnn time series. When computing the variance explained by the design compared to the total variance of the signal (in each respective method dataset – Total SS), denoising resulted in an increasingly higher portion of variance explained by the experimental design (Figure 9A). This is in line with the increased statistical detection sensitivity afforded by NORDIC denoising (with or without the noise scan - Figure 2). Interestingly though, after NORDIC, information related to the experimental design was present in the part of the signal that was removed by the denoising procedure. In relation to the total original variance (Total SS Original), the variance explained by the design in the residuals after NORDIC was higher for NORdef compared to NORnn, which is in line with the higher number of principal components that are removed when using NORdef compared to NORnn. Similar patterns of variance explained in the data or the residuals after NORDIC were visible across all individual subjects (Figure 10). Permutations indicated that the effects described were significant against and alpha level of 0.05, corrected for multiple comparisons, both for the increase in variance explained by the design in the time series (before and after NORDIC) and for the increase in variance explained by the design in the residuals after NORDIC (p<0.001).

**Figure 9.**
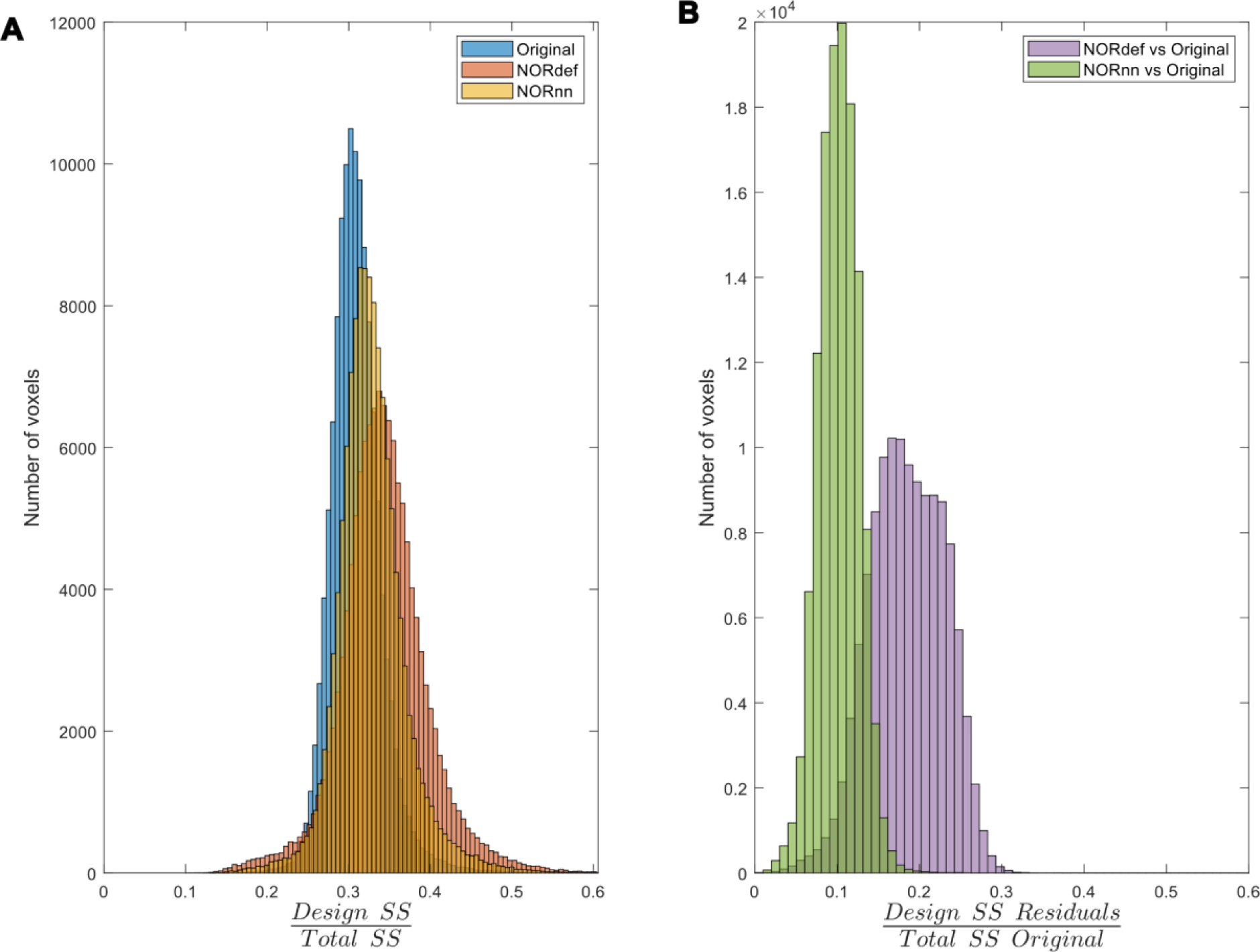
Variance partitioning. A) The amount of variance explained by our design in the data increases consecutively with the use of NORnn and NORdef respectively for one exemplary participant. B) Denoising results in the removal of part of the signal. A proportion of the variance in the residuals after NORDIC can be explained by our stimulation design.

**Figure 10.**
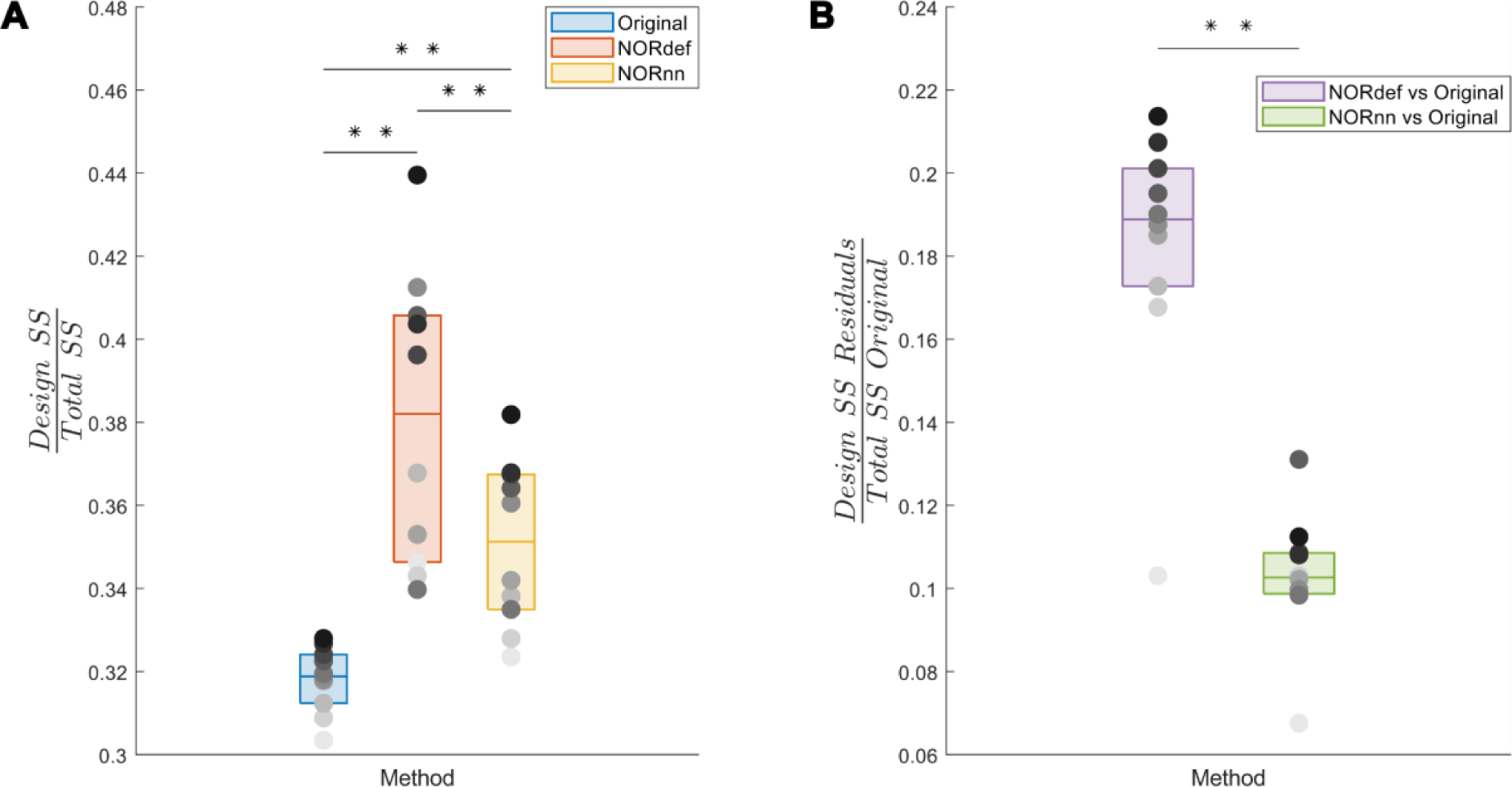
Group analysis of the variance explained by the stimulation design. A) Box charts show the interquartile percentile range of variance explained by the design in the data across participants. After NORDIC denoising, an increased proportion of the variance is explained by the experimental design. NORnn shows an increase in explained variance compared to the original data, but a slightly lower increase than NORdef. B) The proportion of variance explained by the design that is removed from the Original data after NORDIC. NORdef removed a larger proportion of the signal compared to NORnn. * indicates p<0.05, ** indicates p<0.01.

### 3.3 Laminar data

Submillimeter data collected with experimental designs presented here are often used to investigate task-related cortical depth dependent changes in functional activity. In preparation for such future studies, we set out to determine the effect NORDIC has on the laminar profiles. We considered the depth dependent changes (11 equivolume cortical depths) associated with the PredH condition. While in all participants, we could observe the expected increase towards the surface in our GE-BOLD data (Heinzle et al., 2016; Menon et al., 1995; Turner, 2002), NORDIC denoising is associated with a clear reduction in percent signal in superficial cortical depths (see Figure 11 for three representative subjects).

**Figure 11.**
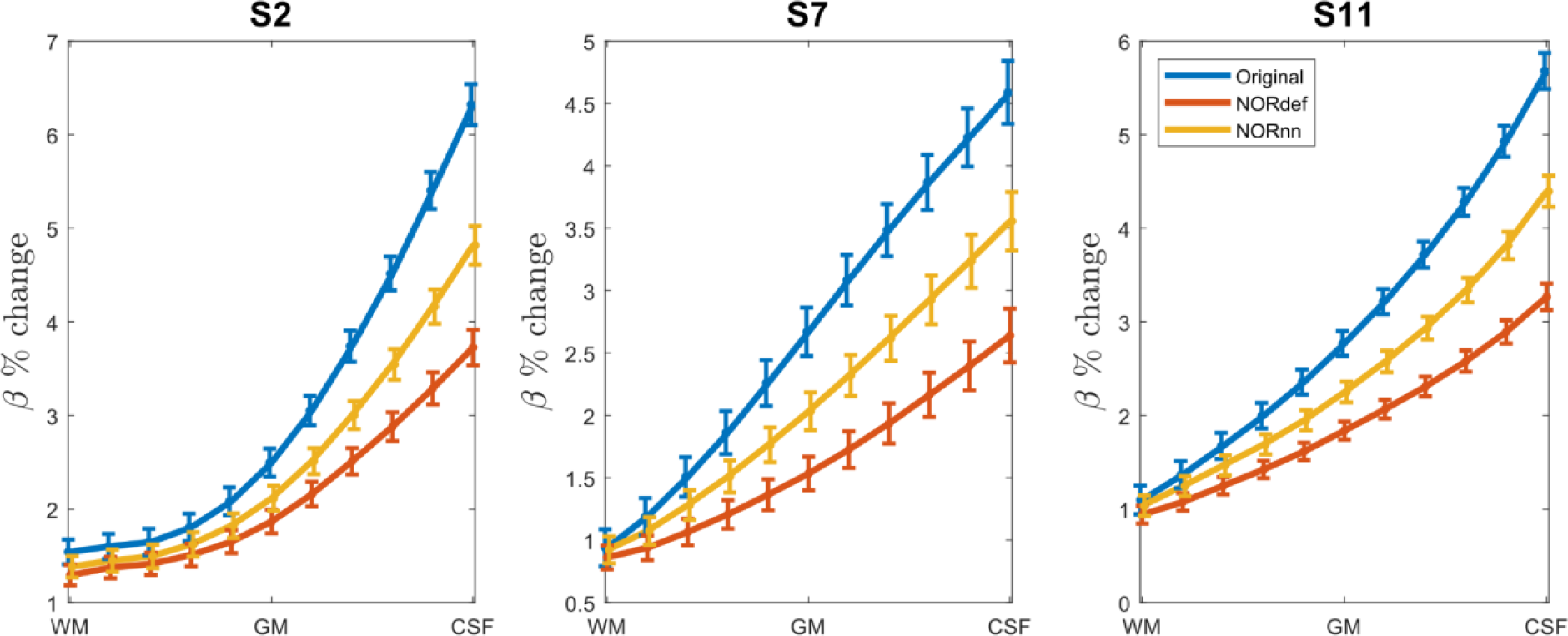
Effect of NORDIC across depth. For three participants we plot the laminar response profiles for the PredH condition. In all plots we can easily identify the draining vein effect. However, we see a gradual decrease in slope for NORnn and NORdef, indicating that NORDIC denoising has a differential effect across depths.

## 4 Discussion

Functional MRI is an indispensable tool for the investigation of human brain function. However, fMRI data is inherently limited by physiological and thermal noise (Triantafyllou et al., 2005, 2011). For this reason, in the fMRI community, the development of methods for removing unwanted sources of variance in the data has been a longstanding goal. Denoising techniques in fMRI can be broadly distinguished in those that tackle the removal of structured (physiological) noise and those that instead aim to reduce thermal noise. A technique that has been introduced to deal with thermal noise in particular is NORDIC PCA. NORDIC denoising has been vetted in various brain areas, voxel sizes, experimental designs and field strengths (for examples see Dowdle et al., 2022, 2023; Knudsen et al., 2023; Raimondo et al., 2023; Vizioli et al., 2021). While in fMRI several approaches have been introduced to improve (statistical) detection power of the signals of interest, it is important to note that any denoising approach may affect the temporal or spatial precision of the underlying fMRI signal as well as their bias-variance tradeoff (Kay, 2022). Ideally, denoising techniques should not spatially or temporally blur the data, while also minimizing any bias introduced. In its initial applications to fMRI, NORDIC denoising has been shown to preserve spatial and temporal information as well as not introducing unwanted biases in the data. These applications have focused primarily on visual and motor cortical areas at different magnetic field strengths (3T and 7T) and using different experimental designs as well as contrast mechanisms (Dowdle et al., 2022, 2023; Knudsen et al., 2023; Raimondo et al., 2023).

Investigating fMRI responses in temporal cortical areas with high spatial resolution (at UHF) is particularly challenging. The location of (primary) cortical areas in particular calls for large field-of-view acquisitions (in either transversally or coronally applied slices to ensure bilateral coverage), which requires high in-plane acceleration to reduce distortions in the resulting EPI images. In addition, when using a single transmit coil, as is the case in most applications, inhomogeneities in the radio frequency transmit field result in suboptimal flip angles (Moerel et al., 2021). While ad-hoc solutions can be found (e.g. by limiting the coverage to single hemispheres), high spatial resolution investigations of temporal cortical areas result in lower temporal SNR compared to e.g. visual or motor cortical regions. For these reasons, we evaluated the consequences associated with the use of NORDIC denoising in temporal cortical areas and extend this to a larger number of subjects. We compared two processing strategies to the original data (i.e. no NORDIC denoising), one dataset using the default settings for fMRI NORDIC (i.e. using magnitude and phase images and a noise threshold estimated using noise scans) and one dataset with a more conservative noise threshold obtained from g-factor estimation.

Our results indicate that NORDIC processing results in increased reliability of the response estimates, increased reliability of the spatial patterns and increase similarity of single run patterns to an ideal model formed by averaging multiple runs. However, our results suggest that, in auditory cortical regions, NORDIC denoising is associated with a non-negligible difference in the percent signal changes, compared to the original data, elicited by our slow event-related design (Figures 6 and 8). These effects are reminiscent of regularization approaches in regression as it results in lower estimated regression coefficients (i.e. betas) while reducing their variance. The variance reduction is proportionally larger with respect to the introduced bias, as evidenced by the increased t-statistics (Figures 6 and 8) and underlies the increased statistical detection sensitivity following NORDIC processing compared to the original data (Figure 2 - and in agreement with previous studies Dowdle et al., 2023; Vizioli et al., 2021). Importantly, the reduced variance in the NORDIC processed data results in increased spatial consistency (especially when evaluated in a repeated split half analysis). All our analyses performed at the level of beta estimates in different temporal cortical regions showed a gradual improvement (e.g. in t-statistics) from NORnn to NORdef (and an associated larger bias in NORdef compared to NORnn), in line with our assumption that NORnn is the more conservative approach. Interestingly, even within a dataset, the deviations from the original data introduced by NORDIC are not uniform, it is associated with the amount of signal present in the data. That is, voxels with more signal (as measured by the mean of the time series divided by the standard deviation of the time series [tSNR]) show the lowest change in estimated percent signal (Figure 7).

At the group level, the lower variance associated with the estimated responses also resulted in a higher run-to-run correlation (with significant effects at the group level observed for NORnn, see Figures 3 and 4). It is interesting to note that when considering the run-to-run variability or the correlation to a multi-run reference, NORDIC seems to improve data in most, but not all of our participants. For participants in which the original data exhibit the lowest reliability the improvements are not noticeable (see single participants points in Figure 4). We can only speculate about the reason for the lack of improvement. These two participants displayed the most movement across their scanning session, which may have resulted in the noise in the data being mainly physiological of origin. This could be a reason why NORDIC denoising did not result in a large improvement for these two participants.

The difference between the original data and NORDIC processed data is suggestive of the fact that some signal (associated with the experimental design) has been removed by the approach. We confirmed this by analyzing the portion of the signal from the magnitude images that is removed by NORDIC (computed as the portion of the original data time series orthogonal to either the NORdef or NORnn time series). While the design explained larger portions of variance in the data after NORDIC processing, the design also explained larger portions of variance in the residuals after NORDIC (Figures 9 and 10). This indicates, that perhaps not surprisingly, NORDIC can remove portions of the signal that in a given sample (i.e. a functional run) are indistinguishable from the noise. These results are in agreement with the results indicating larger changes in beta estimates after NORDIC (compared to the original data) in voxels with lower tSNR (putatively voxels in which the signal and the noise are more confounded - Figure 7).

As a preliminary analysis, we investigated the difference in laminar profiles between NORDIC and the original data (Figure 11). The larger changes in estimated percent signal were noticeable on superficial cortical layers (and more so for NORdef compared to NORnn). This interesting effect may relate to the changes in signal and noise contributions across depths in GE-fMRI. Further research is necessary to explore the causes of these changes induced by NORDIC in the layer dependent signals and the consequences they may have on neuroscientific conclusions drawn by investigating differential responses across layers, or when more elaborate modeling techniques are used (Markuerkiaga et al., 2016; Uludag & Havlicek, 2021; van Mourik et al., 2019).

It is important to note that we here defined the bias introduced by NORDIC as the reduction in percent signal changes that is visible when analyzing the time series after NORDIC compared to the original data (Figures 6 and 8). While NORDIC acts on complex data (to ensure a Gaussian distribution of the noise) the percent signal estimates are computed on the magnitude data. The noise distribution in magnitude only data is not Gaussian but Rician (see e.g. Manzano-Patron et al., 2023) and can result in a biased estimate of the effects. That is, it is possible that the reduced percent signal change we observe after NORDIC is stemming from a larger bias in the estimates obtained from the original data induced by the elevated noise floor. While this explanation offers an alternative interpretation of the reduced percent signal changes obtained after NORDIC, it is not clear how it can explain the effects we report on the portion of the variance explained by the design in the residuals of the time series after NORDIC (Figure 9 and Figure 10). This is because any amplitude difference between the original and the NORDIC time series is accounted for in the way we estimate the residuals after NORDIC (i.e. these residuals are not a simple subtraction of the data before and after NORDIC).

Our results have some implications for the use of NORDIC in neuroscientific investigations as well as for future methodological developments of this denoising technique. First, as NORDIC can (in low SNR regimes as ours) remove portions of the signal, it follows that its application on a run-to-run basis may not combine its benefits to the more general practice of averaging. That is, while averaging will preserve all signal portions in the single run data (and with enough runs may render small effects detectable), NORDIC may remove some of these effects in the single runs and make them undetectable even after extensive averaging. Second, any biases introduced by NORDIC is likely related to signal components that, in a given sample (i.e. a run) are indistinguishable from noise. This consideration highlights the need to further investigate the interaction between the experimental design and any bias introduced by NORDIC processing. That is, in our data the effect may have been exacerbated by the slow event-related stimulus presentation that may confound the response (i.e. the signal) more with the noise in low SNR regimes. While in visual areas event-related designs do not result in a detectable bias after NORDIC (Dowdle et al., 2023), this may relate to the higher SNR of visual areas compared to temporal regions. Finally, it is tempting to speculate that several approaches could be undertaken to abate the bias. Here, we showed that a more conservative threshold for the identification of noisy eigenvalues results in a lower bias (NORnn). Further investigations are warranted in evaluating the effect that other settings (e.g. the patch size) have on the bias. More sophisticated approaches could be considered to, for example, select principal components for removal only if their relationship with the experimental design is negligible akin to the selection of interesting components when performing independent component analysis for task fMRI (De Martino et al., 2007; McKeown et al., 1998; Moritz et al., 2005; Schmithorst & Brown, 2004). Such an approach would not generalize to resting state fMRI but could help for task based functional studies.

Independent of the biases we describe here, NORDIC processing remains an important tool for fMRI investigations especially when SNR is limited (i.e. when thermal noise is dominant), such as laminar studies or functional MRI studies using less sensitive contrast mechanisms (e.g. spin-echo BOLD or non-BOLD contrast mechanisms such as cerebral blood flow-based vascular space occupancy or blood flow based contrast mechanisms such as arterial spin labeling). NORDIC can then be used as a complement to techniques that target physiological noise components to improve the usability of these different SNR starved acquisition approaches. Similarly, NORDIC could be very beneficial in patient studies that cannot rely on long scan times (i.e. extensive averaging) because of practical constraints. In general, though, while any given processing or reconstruction step likely introduces some bias, and while it may be acceptable in some circumstances, it is reasonable to advise NORDIC users to evaluate the amount of bias introduced in their data (by e.g. plotting percent signal estimates before and after NORDIC) apart from focusing only on the increased (statistical) detectability of the effects.

In conclusion, NORDIC can be added to the family of preprocessing techniques that can be utilized to improve the detection sensitivity and reliability of the responses estimated from the fMRI signal. The improvements NORDIC affords warrant its use in SNR challenged settings. Following previous reports, also in our data these positive effects were significant. The signal changes we report here, on the other hand, suggest that some care is required when using NORDIC – new applications may have to further characterize the effect of NORDIC to better evaluate the generalizability of its effects.

## Data and Code Availability

Analyses codes (after preprocessing) are available on Github https://github.com/lonikefaes/auditory_nordic. The anonymized raw data of this study is available and can be downloaded from doi:10.18112/openneuro.ds004928.v1.0.0. (The dataset will be made publicly available upon acceptance).

## Author Contributions

**Lonike K. Faes:** Conceptualization, Formal Analysis, Methodology, Visualization, Writing – Original Draft, Writing – Review & Editing **Agustin Lage-Castellanos:** Conceptualization, Formal Analysis, Methodology, Visualization, Writing – Review & Editing **Giancarlo Valente:** Methodology, Writing – Review & Editing **Zidan Yu:** Investigation, Resources **Martijn A. Cloos:** Investigation, Resources, Writing – Review & Editing **Luca Vizioli:** Investigation, Resources, Writing – Review & Editing **Steen Moeller:** Writing – Review & Editing **Essa Yacoub:** Conceptualization, Funding Acquisition, Writing – Review & Editing **Federico De Martino:** Conceptualization, Funding Acquisition, Methodology, Supervision, Writing – Original Draft, Writing – Review & Editing.

## Funding

The research was supported by the National Institute of Health (RF1MH116978-01 and P41EB027061). Additionally, this research was supported by the European Research Council (ERC) under the European Union’s Horizon 2020 research and innovation program (grant agreement No. 101001270) awarded to FDM.

## Declaration of Competing Interests

The authors declare no conflict of interest.

## Acknowledgements

We would like to thank Logan Dowdle for the useful discussions. Moreover, we would like to thank Renzo Huber and Lasse Knudsen for the helpful discussions and comments on the first draft of this manuscript.

## Notes

### Competing Interest Statement

The authors have declared no competing interest.

