## Supplementary Table 1 for "Evaluating the effect of denoising submillimeter auditory fMRI data with NORDIC"

### Supplementary Material

**Table 1.**

P-values originating from all comparisons depicted in Figure 8. Correction for multiple comparisons using Bonferroni resulted in the lowest possible p-value being 0.015. Similarly, due to the Bonferroni correction, the p-value of certain comparisons is artificially higher than 1 (set to 1.000 in the table).

|  | **Original - NORdef** | **NORdef - NORnn** | **Original - NORnn** |
| --- | --- | --- | --- |
| Beta HG | 0.015 | 0.015 | 0.015 |
| Beta PP | 0.015 | 0.015 | 0.015 |
| Beta PT | 0.015 | 0.015 | 0.015 |
| Beta aSTG | 1.000 | 1.000 | 1.000 |
| Beta pSTG | 0.015 | 0.044 | 0.015 |
| t-value HG | 0.015 | 0.015 | 0.015 |
| t-value PP | 0.015 | 0.132 | 0.015 |
| t-value PT | 0.015 | 0.044 | 0.015 |
| t-value aSTG | 0.015 | 0.015 | 0.015 |
| t-value pSTG | 0.015 | 0.015 | 0.015 |
| Beta stab. HG | 0.044 | 0.220 | 0.015 |
| Beta stab. PP | 1.000 | 1.000 | 1.000 |
| Beta stab. PT | 0.044 | 0.044 | 0.015 |
| Beta stab. aSTG | 1.000 | 1.000 | 0.952 |
| Beta stab. pSTG | 0.718 | 1.000 | 0.103 |
| t-value stab. HG | 0.776 | 1.000 | 0.015 |
| t-value stab. PP | 0.015 | 0.103 | 0.015 |
| t-value stab. PT | 0.015 | 0.044 | 1.000 |
| t-value stab. aSTG | 0.015 | 0.015 | 0.044 |
| t-value stab. pSTG | 0.015 | 0.015 | 0.015 |
